# Anaerobic biodegradation of chloroform and dichloromethane with a *Dehalobacter* enrichment culture

**DOI:** 10.1101/2021.07.26.453918

**Authors:** Hao Wang, Rong Yu, Jennifer Webb, Peter Dollar, David L. Freedman

## Abstract

Chloroform (CF) and dichloromethane (DCM) are among the more commonly identified chlorinated aliphatic compounds found in contaminated soil and groundwater. Complete dechlorination of CF has been reported under anaerobic conditions by microbes that respire CF to DCM and others that biodegrade DCM. The objectives of this study were to ascertain if a commercially available bioaugmentation enrichment culture (KB-1^®^ Plus) uses an oxidative or fermentative pathway for biodegradation of DCM; and to determine if the products from DCM biodegradation can support organohalide respiration of CF to DCM in the absence of an exogenous electron donor. In various treatments with the KB-1^®^ Plus culture to which ^14^C-CF was added, the predominant product was ^14^CO_2_, indicating that oxidation is the predominant pathway for DCM. Recovery of ^14^C-DCM when biodegradation was still in progress confirmed that CF first undergoes reductive dechlorination to DCM. ^14^C-labeled organic acids, including acetate and propionate, were also recovered, suggesting that synthesis of organic acids provides a sink for the electron equivalents from oxidation of DCM. When the culture was washed to remove organic acids from prior additions of exogenous electron donor and only CF and DCM were added, the culture completely dechlorinated CF. The total amount of DCM added was not sufficient to provide the electron equivalents needed to reduce CF to DCM. Thus, the additional reducing power came via the DCM generated from CF reduction. Nevertheless, the rate of CF consumption was considerably slower in comparison to treatments that received an exogenous electron donor.

**IMPORTANCE:** Chloroform (CF) and dichloromethane (DCM) are among the more commonly identified chlorinated aliphatic compounds found in contaminated soil and groundwater. One way to address this problem is to add microbes to the subsurface that can biodegrade these compounds. While microbes are known that can accomplish this task, less is known about the pathways used under anaerobic conditions. Some use an oxidative pathway, resulting mainly in carbon dioxide. Others use a fermentative pathway, resulting in formation of organic acids. In this study, a commercially available bioaugmentation enrichment culture (KB-1^®^ Plus) was evaluated using carbon-14 labelled chloroform. The main product formed was carbon dioxide, indicating the use of an oxidative pathway. The reducing power gained from oxidation was shown to support reductive dechlorination of CF to DCM. The results demonstrate the potential to achieve full dechlorination of CF and DCM to nonhazardous products that are difficult to identify in the field.

## INTRODUCTION

Considerable progress has been made in identifying the microbes and pathways involved in anaerobic biodegradation of chloroform (CF) and dichloromethane (DCM). For many decades, reductive dechlorination of CF to DCM was thought to be a cometabolic process (1). A key discovery by Grostern et al. (2) demonstrated the use of CF as a terminal electron acceptor during organohalide respiration by a *Dehalobacter* population. This built on the an earlier discovery that strain TCA1, a close relative to *Dehalobacter restrictus*, is able to respire using 1,1,1-trichloroethane as its terminal electron acceptor (3). CF and 1,1,1-trichloroethane share the same trichloro structure on a single carbon. The processes are also similar in that reductive dechlorination stops before complete dechlorination is achieved, unlike what is possible with organohalide respiration of chlorinated ethenes (4). With CF, DCM is the main product, while respiration of 1,1,1-TCA, stops at chloroethane, both of which have potential environmental and toxicological impacts. Consequently, to achieve acceptable anaerobic remediation, further dechlorination is required to non-toxic end-products.

Anaerobic biodegradation of DCM as a sole source of carbon and energy has been known for several decades (5, 6). Following development of highly enriched cultures that grow on DCM (7), Leisinger and colleagues isolated *Dehalobacterium formicoaceticum* gen. nov. sp. nov. (8). They subsequently proposed a modified version of the Wood-Ljungdahl pathway for fermentation of DCM to a mixture of formate and acetate, involving a Co(I) corrinoid and tetrahydrofolate (9). More than a decade passed before the next wave of research appeared, with a focus this time on characterization of the RM consortium (10-12) and DCMF enrichment culture (13). DCM degradation by the RM consortium, originally derived from pristine river sediment, was initially attributed to *Dehalobacter* sp. (14, 15). The microbe in the RM consortium responsible for DCM degradation was subsequently identified as *Candidatus* Dichloromethanomonas elyunquensis. The microbe enriched from organochlorine-contaminated groundwater near Botany Bay, Australia that was responsible for DCM degradation was initially identified as a *Dehalobacter* (16). Upon further enrichment, the DCMF culture became dominated by a novel member of the *Peptococcaceae* family.

Various methods have been used to gain insight into the DCM biodegradation pathways used by the RM consortium and DCMF. Strain RM uses a mineralization pathway that oxidizes the carbon in DCM to CO_2_, with the electron equivalents (4 eeq/mol DCM) released as H_2_. The proposed pathway for DCMF is more similar to the fermentation steps used by *Dehalobacterium formicoaceticum*, with the principal products being acetate and formate. Based on free energy calculations, Chen et al. (17) predict that the mineralization pathway is predominant in environments with H_2_ levels below 100 parts per million by volume (ppmv).

Several studies have evaluated concurrent anaerobic biodegradation of CF and DCM. Lee et al. (18) describe a microbial community that respires CF to DCM using emulsified vegetable oil or acetate as electron donors, followed by dehalofermentation of the DCM. Both processes were attributed to *Dehalobacter*. Justica-Leon et al. (14) demonstrated reduction of CF to DCM followed by consumption of DCM in microcosms bioaugmented with the CF-to-DCM organohalide-respiring culture Dhb-CF and the DCM-degrading consortium RM. Electron donor was supplied in the form of lactate and hydrogen. Neither of these studies explored the possibility that complete dechlorination of CF is possible without addition of an electron donor, based on the potential for fermentation products from DCM serving that purpose. A system similar to this has been described for chlorobenzene. Liang et al. (19) combined a culture that respires chlorobenzene to benzene with another culture that ferments the benzene to acetate and hydrogen, which serve as electron donors for the reduction process and therein remove the need for an exogenous electron donor.

KB-1^®^ Plus is a commercially available bioaugmentation culture that is used to detoxify contaminant plumes containing a variety of chlorinated aliphatic compounds. Prior studies have demonstrated its effectiveness in dechlorinating chlorinated ethanes and CF (2, 20, 21). The KB-1^®^ Plus formulation used in this study is a mixed culture that is maintained on CF and is known to contain *Dehalobacter* spp., although the pathway used for biodegradation of DCM formed from CF has not been reported. The objectives of this study were to ascertain if KB-1^®^ Plus uses an oxidative or fermentative pathway for DCM, based on products formed from ^14^C-labeled CF; and to determine if the products from DCM biodegradation can support organohalide respiration of CF to DCM in the absence of an exogenous electron donor.

## RESULTS

### Analysis of CF biodegradation using ^14^C-CF

VOC results for one of the triplicates of treatment #1 (without BES added) and #3 (with BES added) are shown in Figure 1; results for other replicates and repeat experiments, as well as treatments #2 and #4, are shown in Supplementary Materials (Figures S1-S10). Treatment #2 behaved similarly to treatment #1 except that the incubation was stopped as soon as CF was consumed (Figures S4-S6). Only a minor amount of CM was detected in any of the bottles (0.13 μmol maximum, data points not shown), indicating that hydrogenolysis of DCM to CM was not an important part of the CF biodegradation pathway. In the absence of BES, the rate of CF and DCM biodegradation was approximately twice as fast. In both treatments, the maximum accumulation of DCM was approximately one third of the stoichiometric amount of CF, indicating DCM fermentation occurred during hydrogenolysis of CF to DCM. Methanogenesis was inhibited by the presence of CF in treatment #1; once the CF was consumed, an average of 7 µmol of methane accumulated. By contrast, the presence of BES inhibited methanogenesis throughout the incubation. There was only a minor decrease in CF in the medium A control (treatment #4; Figure S10) and no accumulation of DCM, CM, or methane.

**Figure 1.**
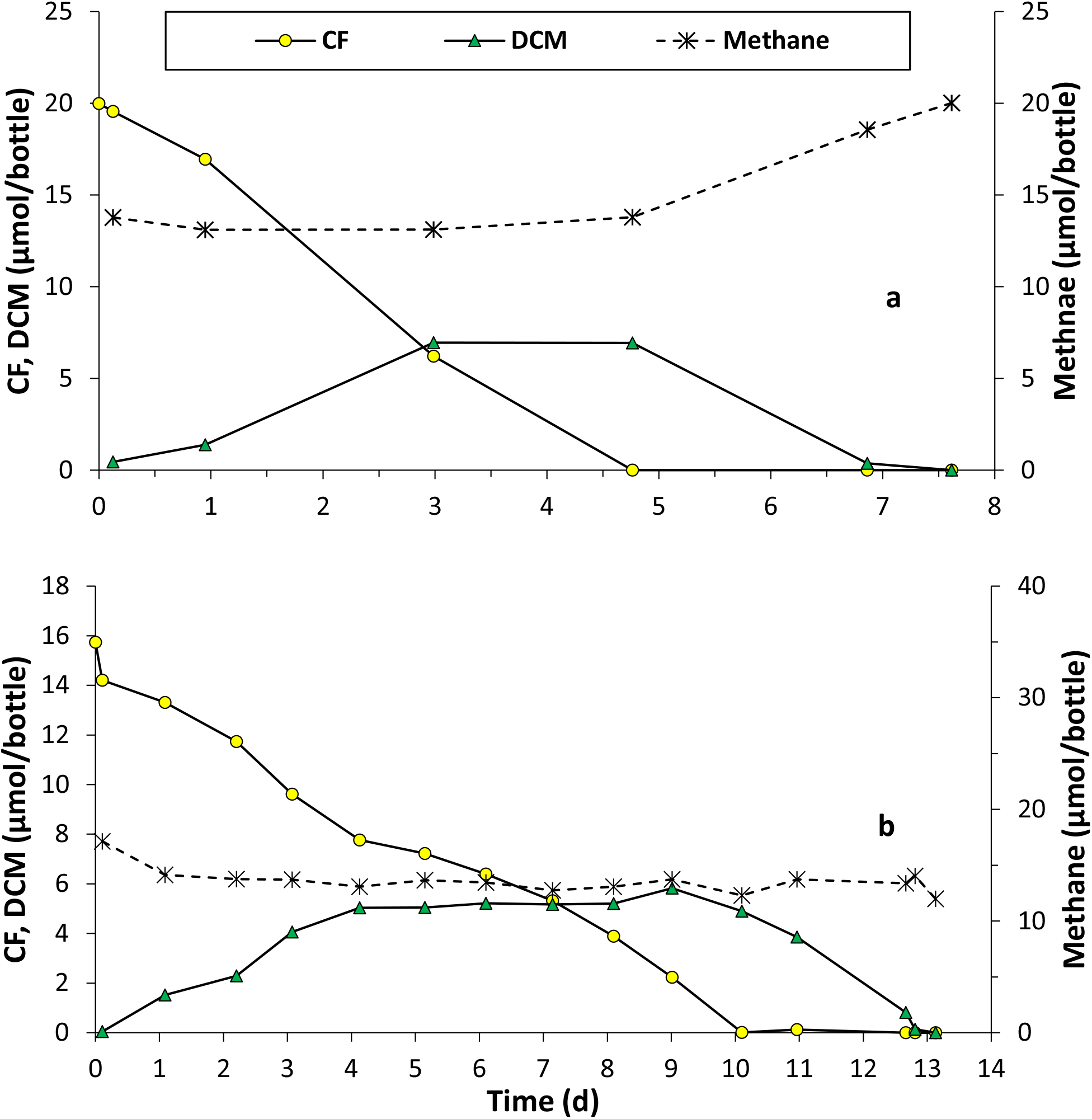
Representative results for biodegradation of CF by the KB-1 Plus^®^ culture with the addition of lactate (electron donor) **a**) treatment #1, without BES added; and **b**) treatment #3, with BES added. One dose of lactate was added to all bottles at time zero. Results for single bottles are shown; replicates behaved similarly (Supplementary Material). Results for treatments #2 and #4 are provided in the Supplementary Material.

The distribution of ^14^C for treatments #1-4 is summarized in Figure 2. The results represent averages for 3 to 9 bottles in each treatment (Table 1). ^14^CO_2_ was the predominant product in treatments with KB-1^®^ Plus present (59-78% of the initial ^14^C-CF added). ^14^C-DCM was significant only in treatment #2 bottles that were sacrificed after CF was consumed but before all of the DCM was consumed. It is notable that there was also significant accumulation of ^14^CO_2_ in these bottles, even before the DCM was completely consumed. ^14^C-NSR, which includes organic acids, comprised an average of 5.3% of the ^14^C-CF added, suggesting that the principal anaerobic biodegradation pathway for CF by KB-1^®^ Plus is via mineralization, rather than direct incorporation of carbon from DCM into organic acids. Samples of NSR from treatment #3 were separated by HPLC. The fraction with the highest level of ^14^C activity (62-69% of the ^14^C in the NSR) had the same retention time as acetate; the next highest fraction (6.4-13% of the ^14^C in the NSR) eluted at the same time as propionate.

**Table 1.** Experimental design.

| Treatment<br># | Contents | n<sup>a</sup> | <sup>14</sup>C-CF<br>(μCi)<sup>b</sup> | CF<br>(μmol)<sup>b</sup> | DCM<br>(μmol)<sup>b</sup> | DCM<br>status<sup>c</sup> | Lactate<sup>d</sup> | Acetate<sup>e</sup> | BES |
| --- | --- | --- | --- | --- | --- | --- | --- | --- | --- |
| 1 | KB-1<sup>®</sup> Plus | 9 (6) | 0.36 | 17.2 | 0 | C | Yes | Yes | No |
| 2 | KB-1<sup>®</sup> Plus | 9 (6) | 0.34 | 17.2 | 0 | NC | Yes | Yes | No |
| 3 | KB-1<sup>®</sup> Plus | 6 (6) | 0.37 | 17.2 | 0 | C | Yes | Yes | Yes |
| 4 | Medium A<sup>f</sup> | 3 | 0.38 | 15.7 | 0 | - | - | - | - |
| 5 | KB-1<sup>®</sup> Plus | 3 | 0 | 0 | 16.5 | C | No | Yes | No |
| 6 | KB-1<sup>®</sup> Plus | 3 | 0 | 0 | 16.5 | C | No | Yes | Yes |
| 7 | KB-1<sup>®</sup> Plus | 3 | 0 | 1.68 | 16.5 | C | No | Yes | No |
| 8 | KB-1<sup>®</sup> Plus/w<sup>g</sup> | 3 | 0 | 4.02 | 0.47 | C | No | No | No |
| 9 | KB-1<sup>®</sup> Plus/w | 3 | 0 | 0 | 0 | - | No | No | No |
| 10 | Medium B<sup>f</sup> | 3 | 0 | 4.02 | 0.94 | - | - | - | - |

<sup>a</sup> Number of microcosms. Values in parenthesis represent number of bottles that received a dose of unlabeled CF prior to adding <sup>14</sup>C-CF.
<sup>b</sup> Initial amount added.
<sup>c</sup> C = complete DCM degradation; NC = DCM degradation was not complete.
<sup>d</sup> Lactate added as the electron donor.
<sup>e</sup> Acetate present at the start of the incubation as a byproduct from prior addition of lactate.
<sup>f</sup> Media A and B are described in Materials and Methods.
<sup>g</sup> /w indicates the culture was washed prior to incubation.

**Figure 2.**
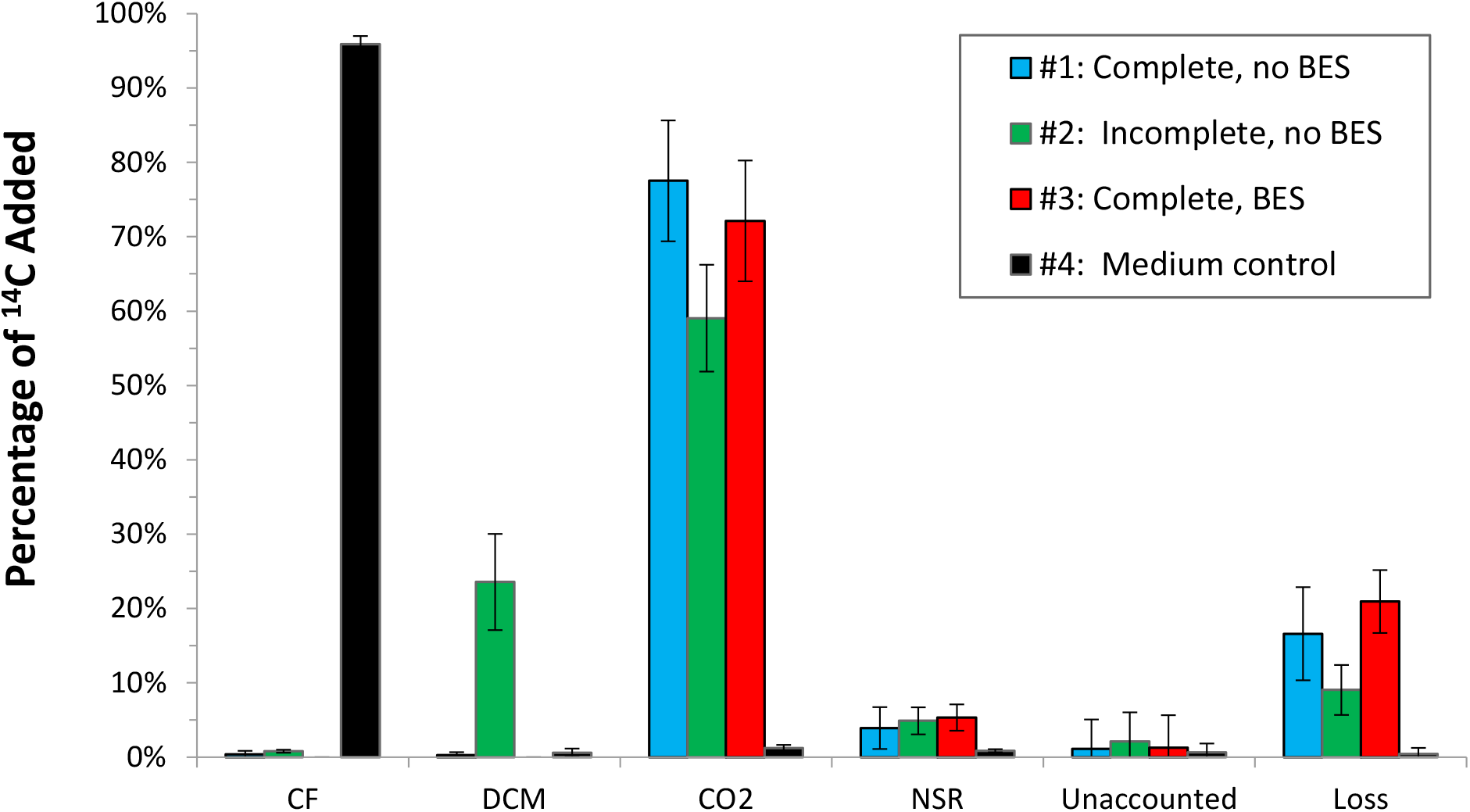
Percentage of ^14^C from ^14^C-CF during biodegradation by KB-1® Plus in treatments #1-4. Error bars represent the standard deviation; the number of replicates for each treatment are shown in Table 1. “Complete” means all of the CF and DCM were consumed. “Incomplete” means the CF was consumed but not all of the DCM was consumed. BES is 2-bromoethanesulfonate, an inhibitor of methanogenesis. Medium A was used for treatment #4.

Nearly complete recovery of ^14^C-CF (96%) and only minor amounts of ^14^C-NSR and ^14^CO_2_ in the medium control confirmed the high purity of the ^14^C-CF added and the lack of abiotic transformation (Figure 2). The amount of ^14^C that was not accounted for was minor. The extent of losses was a function of the incubation time, ranging from an average of 0.46% for the medium control (7 days of incubation) to 21% for treatment #3 (23 days of incubation).

### Evaluation of CF and DCM as sole substrates

When DCM was added to the KB-1^®^ Plus culture without CF (treatment #5), the initial dose (15.6 µmol) was consumed within 3 days, with no accumulation of H_2_ and methane formation that exceeded the level expected from DCM alone, i.e., 0.5 mol CH_4_/mol DCM (Figure 3a). When the DCM dose was increased by an order of magnitude, a lag of approximately 30 days was followed by a rate of DCM consumption similar to the initial dose. The amount of methane produced during consumption of the second dose of DCM was closer to stoichiometric (i.e., 0.41 µmol CH_4_/µmol DCM), and was also accompanied by an increase in hydrogen. However, hydrogen represented a lower percentage of the electron flow versus methane (i.e., 6 µmol of H_2_ accounts for 3 µmol of DCM, out of ∼180 µmol consumed).

**Figure 3.**
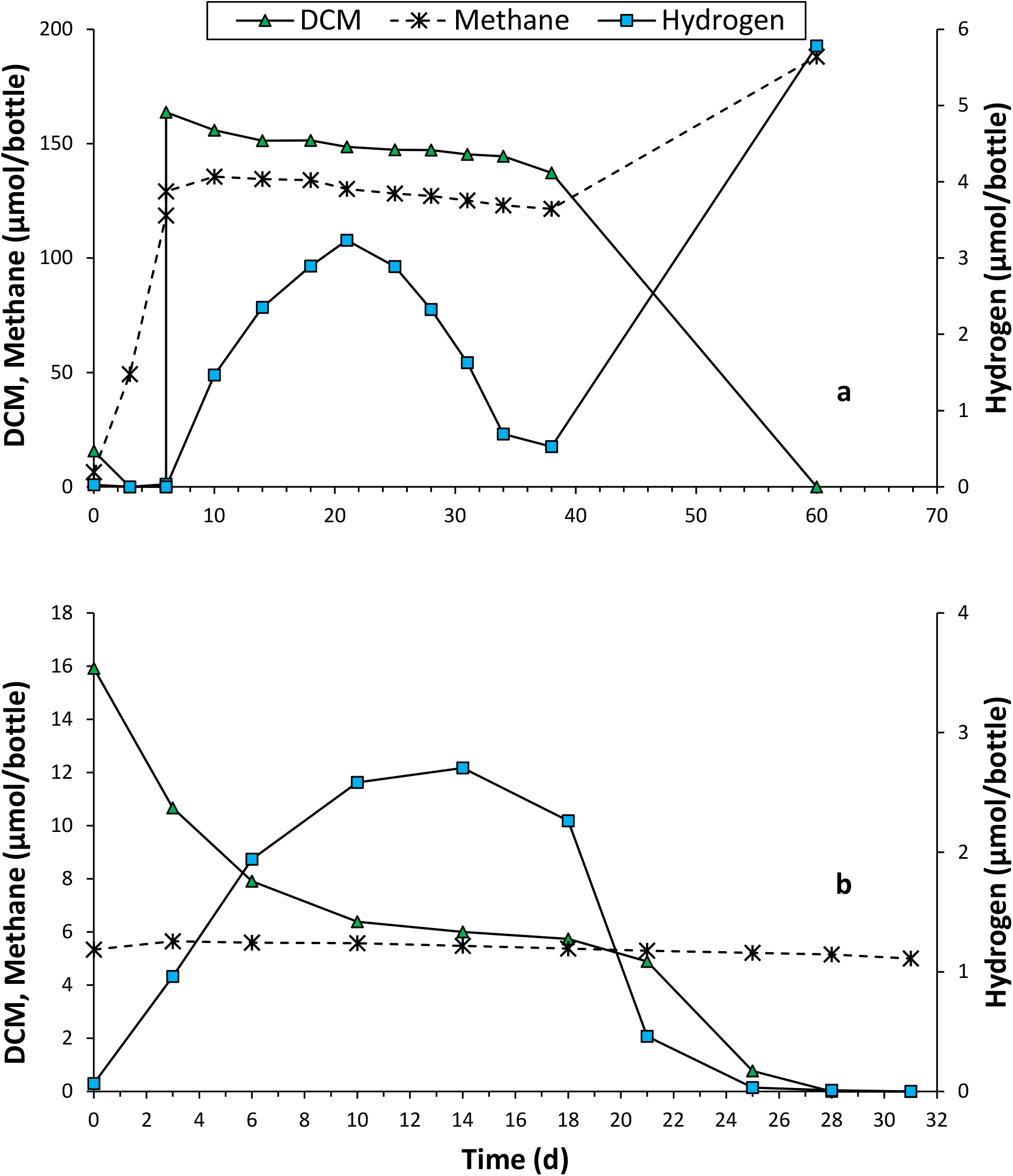
Biodegradation of DCM by KB-1® Plus **a**) without BES added (treatment #5); and **b**) with BES added (treatment #6). Results for single bottles are shown; replicates behaved similarly (Supplementary Material).

When BES was present along with the DCM in treatment #6, the rate of DCM consumption was slowed and there was no accumulation of methane (Figure 3b). A transient decrease in the rate of DCM consumption occurred when hydrogen peaked at approximately 3 µmol per bottle; as H_2_ levels declined, a higher rate of DCM consumption resumed. For treatments #5, and #6, CM remained below 0.5 µmol per bottle.

The observation of hydrogen accumulation during consumption of DCM suggests that DCM may serve as the electron donor for reduction of CF to DCM. This was evaluated with treatment #7, using the KB-1^®^ Plus culture as received. Consequently, acetate was initially present (as a product of lactate added as the electron donor). Repeat additions of ∼17 µmol of CF and DCM per bottle were consumed within several days (Figure 4a), accompanied by accumulation of methane and no detectable hydrogen. As the dose of CF was increased around day 60, methane accumulation stopped, and DCM increased as CF was consumed. Once CF was below detection, the DCM was then completely consumed. This is best seen when the highest dose of CF was added on day 132. DCM increased to 75 µmol per bottle on day 306 and then decreased below detection by day 336, accompanied by an increase in H_2_ of 4.1 µmol per bottle. The next three doses of CF and DCM followed a similar pattern.

**Figure 4.**
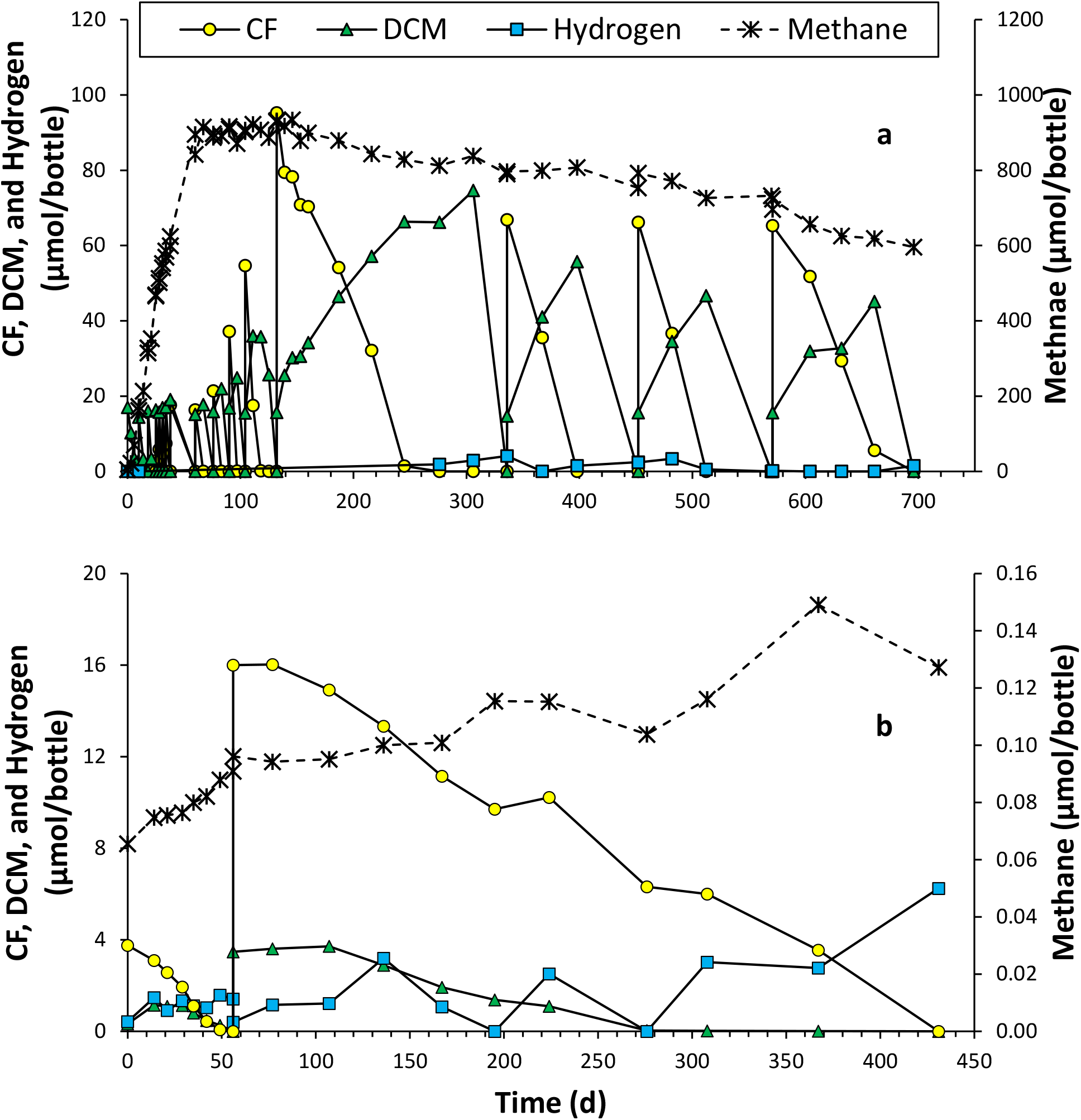
Biodegradation of CF by KB-1® Plus without the addition of sodium lactate in **a**) treatment #7 and **b**) treatment #8. DCM was added to treatment #7 on days 0, 10,18, 25, 28, 31, 34, 38, 60, 76, 90, 132, 336, 452, and 571. DCM was added to treatment #8 on days 0, 14, and 56 (0.33, 0.16, and 3.47 µmol per bottle, respectively). Results for single bottles are shown; replicates behaved similarly (Supplementary Material).

To remove the potential impact of acetate serving as an electron donor for CF reduction to DCM, one of the bottles from treatment #7 was washed and resuspended in medium B to create the inoculum used for treatment #8. The initial dose of CF (3.8 µmol/bottle) and two doses of DCM (0.33 and 0.16 µmol per bottle on days 0 and 14, respectively) were consumed in 56 days (Figure 4b). The increase in methane over this interval (0.03 µmol/bottle) was minor relative to the amount possible (i.e., 0.5 µmol CH_4_ per µmol DCM) based on the peak level of DCM. A second dose of CF (16 µmol/bottle) on day 56 was accompanied by DCM (3.6 µmol/bottle), and DCM was consumed 155 days before the CF. At the end of the incubation period, hydrogen accumulated to 6.3 µmol per bottle. Aqueous samples from treatments #8, 9 and 10 were evaluated for organic acids at the start and end of the incubation. Although small peaks were evident in the HPLC chromatograms at the end of the incubation period, none of the organic acids exceeded the detection limit in treatments #8 and #10. For treatment #9, formate and acetate were slightly above the detection limit (36 and 26 µM, respectively), perhaps from decay of the inoculum. Lactate, propionate, and citrate were below detection in all of the bottles.

## DISCUSSION

Based on the ^14^C results (Fig. 2), a proposed pathway for anaerobic biodegradation of CF by the KB-1^®^ Plus culture is shown in Figure 5. ^14^CO_2_ was the predominant product for the live treatments (#1, #2, and #3). With treatment #2, the bottles were sacrificed when the CF was consumed but DCM was still present. As expected, the DCM was ^14^C-labelled, confirming that DCM is formed via hydrogenolysis of CF. Accumulation of ^14^CO_2_ indicates that the KB-1^®^ Plus culture uses an oxidative pathway for anaerobic biodegradation of DCM, similar to the RM consortium (17). The medium used for cultivation contains 20 mM of bicarbonate. If all 17.2 µmol of CF per bottle was oxidized (i.e., ignoring cell synthesis), the resulting concentration of CO_2_ would be 0.17 mM. Because this is orders of magnitude lower than the bicarbonate level in the medium, the ^14^CO_2_ formed was diluted by a much larger pool of bicarbonate, which would predominate over CO_2_ in the circumneutral medium.

**Figure 5.**
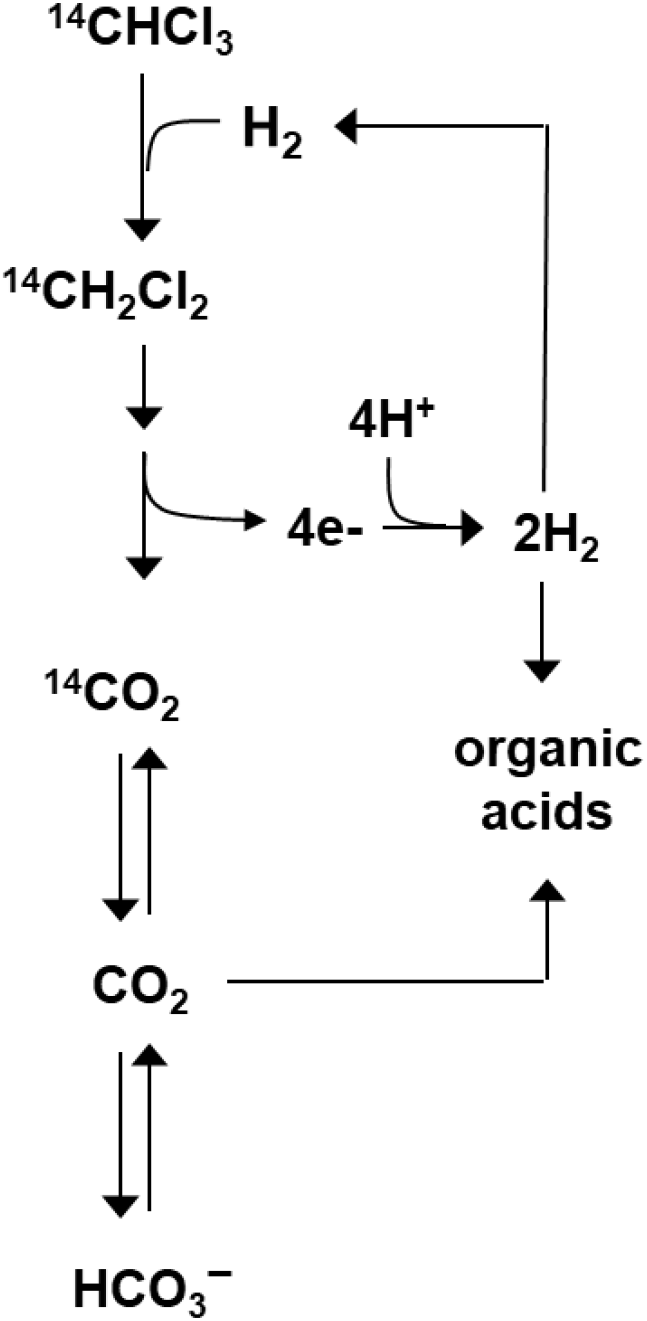
Proposed pathway for anaerobic biodegradation of CF and DCM by the KB-1® Plus culture.

Oxidation of DCM to CO_2_ yields 4 electron equivalents (eeq) per mol, or 2 mol H_2_ (Fig. 5). One mol H_2_ is then available to accomplish reduction of 1 mol CF to DCM. That leaves a net of 1 mol H_2_ per mol of CF. With treatments #5-8, H_2_ accumulated, but even the highest levels observed were never close to the stoichiometric level (Figs. 3 and 4). That indicated there must be a sink for this reducing power. Formation of organic acids is one possibility. With treatments #1-3, approximately 3-5% of the ^14^C was recovered as nonstrippable residue (NSR), and HPLC analysis of NSR from treatment #3 indicated the main components were (presumptively) acetate and propionate. However, because the ^14^CO_2_ generated from ^14^C-CF was diluted in bicarbonate present in the medium, most of the organic acids formed would likely be unlabeled. This would make them difficult to quantify in treatments #1-3, because of the background level of organic acids present in the KB-1^®^ Plus culture as received from SiREM.

One of the objectives with treatment #8 was to improve the potential to quantify the amount of organic acids that can form when CF is biodegraded by the KB-1^®^ Plus culture. The culture in these bottles was washed to remove the background level of organic acids prior to addition of CF and lesser amounts of DCM (Fig. 4). If all of the 19.8 µmol of CF and 4.0 µmol of DCM were oxidized to CO_2_, then 55.6 eeq of reducing power would be available to form organic acids (i.e., 19.8 µmol CF*2 µeeq/µmol + 4.0 µmol DCM*4 µeeq/µmol). Conversion of 55.6 eeq per bottle (with 44 mL/bottle) to acetate (8 µeeq/µmol) would yield a maximum concentration of 158 µM. If the reducing power was used to make formate (6 µeeq/µmol) the outcome would be 211 µM. These concentrations are well above the detection limit. Nevertheless, at the end of the incubation period, there was no statistically significant accumulation of organic acids in comparison to the controls. The extended incubation period (431-477 days) may have complicated closing the mass balance on electron equivalents. If the KB-1^®^ Plus culture can be acclimated to more rapid consumption of CF and DCM in the absence of an exogenous substrate, it may become more feasible to close the balance. Another approach would be to substantially lower the bicarbonate concentration in the medium, so that more of the ^14^CO_2_ could be used to form organic acids.

Another potential sink for the eeq from oxidation of DCM to CO_2_ is methanogenesis. The maximum potential yield is 0.25 mol CH_4_/mol CF and 0.50 mol CH_4_/mol DCM. Having a high background level of organic acids in treatments #1-3 and #5-7 limited the option to quantify methane formation that is attributable only to CF and DCM (e.g., Fig. 1a, 3a, and 4a). With removal of the residual organic acids from treatment #8, the maximum possible yield of methane from the CF and DCM consumed was 7.0 µmol per bottle. The observed increase in the treatment #8 bottles was less than 0.10 µmol per bottle, well below what is needed to close the eeq balance. Furthermore, with treatments in which methanogenesis was inhibited (i.e., #3 and #6 with BES added, and #7 when the CF concentration was increased above ∼22 µmol/bottle, or an aqueous phase concentration of ∼24 mg/L), CF was still consumed to completion, so that methanogenesis was eliminated as a sink for the eeq from oxidation of DCM.

Several studies have reported that oxidation of 1 mol of DCM to 1 mol of CO_2_, 2 mol of H_2_, and 2 mol of Cl^-^ is thermodynamically favorable even at a hydrogen partial pressure of 1 atm (5, 17). This is made possible by the large Gibbs free energy change derived from breaking the carbon chlorine bonds. In spite of the thermodynamic favorability of DCM oxidation even at 1 atm H_2_, the RM consortium requires the presence of H_2_-consuming partners. It is evident from this study that lower levels of H_2_ also favor higher rates of CF consumption by the KB-1^®^ Plus culture (Fig. 3). Inhibition of methanogenesis (with either BES or higher levels of CF) led to slower rates of CF and DCM degradation (Fig. 1, 3, and 4). Methanogenesis contributes to keeping the partial pressure of hydrogen at a low level. Chen et al. (17) speculated that H_2_ inhibition may be related to negative impacts on H_2_-evolving hydrogenases present in the RM consortium.

For treatments #1-3 and #5-7, accumulation of DCM was never stoichiometric with respect to the amount of CF added, indicating that DCM biodegradation was underway as CF was reduced to DCM. Nevertheless, DCM typically increased as CF was consumed and a decreasing trend for DCM did not occur until CF was mostly or completely consumed (Fig. 1, 4a). This is consistent with a previous report that CF inhibits DCM biodegradation, even at CF concentrations as low as 5 mg/L (15). The fact that repeat additions of CF and DCM were consumed indicates that this inhibition is reversible with the KB-1^®^ Plus culture.

The results from treatment #8 support the hypothesis that complete dechlorination of CF is possible without addition of an exogenous electron donor. The source of electron donor for CF reduction to DCM is the oxidation of DCM. The culture used for treatment #8 was washed to remove the accumulated organic acids, yet biodegradation of CF proceeded. The process was “jump started” by adding DCM along with both doses of CF (Fig. 4). Nevertheless, the total amount of DCM added (3.96 µmol/bottle) provided 15.8 µeeq of reducing power, while the total amount of CF consumed (19.75 µmol/bottle) required 39.5 µeeq of reducing power. Thus, the addition reducing power needed must have been made available by the DCM generated from CF reduction. While sustaining CF dechlorination without an input of electron donor is of interest, the rate of CF consumption was considerably slower in comparison to treatments that received electron donor in the form of lactate (Fig. 1). For this reason, it is still advisable to use an electron donor in the field to accelerate the rate of CF reduction to DCM.

The results of this study confirm that the KB-1^®^ Plus culture completely dechlorinates CF to benign products via oxidation of DCM. Documentation of the degradation pathway is especially useful for bioremediation because the products formed are not discernable from the background, so it is not possible to use product accumulation as evidence of in situ biodegradation. Quantifying the presence of the necessary microbes for CF reduction and DCM biodegradation, combined with other lines of evidence such as compound specific isotope analysis (29, 30), provide robust opportunities to document in situ bioremediation of CF.

## MATERIALS AND METHODS

### Chemicals and enrichment culture

CF (99.7%) was obtained from Shelton Scientific. Methane (99.5%), H_2_ (99.995%), N_2_ (99.998%), chloromethane (CM, 99.9%), and dry breathing air were obtained from Airgas. DCM (99.99%) was obtained from Burdick & Jackson; sodium lactate syrup (containing 58.8-61.2% sodium lactate; specific gravity = 1.31) from Spectrum Chemical MFG Corporation; sodium formate (99.3%), sodium citrate dihydrate (>99%), and resazurin (sodium derivative) from J.T. Baker; sodium acetate (>99%) from EM Science; ^14^C-CF from American Radiolabeled Chemicals (0.5 mCi/mmol); Scintisafe Plus liquid scintillation cocktail (LSC) (50%) from Fisher Scientific; and 2-bromoethanesulfonic acid (BES; sodium salt, 98%) and sodium propionate (99%) from Sigma Aldrich. All other chemicals were reagent grade. Medium A was used as a control to assess abiotic losses of ^14^C-CF (see below); its composition is described previously (22). The KB-1^®^ Plus enrichment culture and lactate-free anaerobic mineral medium B were provided by SiREM and shipped on ice to Clemson University via an overnight carrier. Upon receipt, the culture and the medium were placed in an anaerobic chamber and allowed to warm to room temperature.

### Experimental design

The experimental design includes the use of ^14^C-CF, the presence of various electron donors, and use of BES to inhibit methanogenesis (Table 1). One set of experiments (treatments #1-4) was performed with ^14^C-CF for the purpose of determining the ^14^C products that formed. These experiments utilized KB-1^®^ Plus (CF formulation) as received and lactate as the primary electron donor. A second set of experiments evaluated use of DCM as an electron donor in support of hydrogenolysis of CF to DCM. Treatments #5 and 6 evaluated DCM as the sole substrate added; treatments #7-9 included both CF and DCM. Treatment #10 served as a control for product formation from the inoculum used in treatment #8. Acetate was present as a fermentation product from lactate except when it was removed by washing the culture (treatments #8 and 9; described below).

Treatments #1-7 were prepared in 160 mL serum bottles with 100 mL of culture or medium; treatments #8 and 9 were prepared in 70 mL serum bottles with 44 mL of washed culture or medium; treatment #9 was prepared in 12 mL serum bottles with 3 mL of washed culture. The serum bottles were capped with Teflon-faced grey butyl rubber septa; the bottles and septa were autoclaved prior to use. After dispensing the culture in an anaerobic chamber, the serum bottles were removed, the headspace was sparged with N_2_/CO_2_ (70%/30%) to remove hydrogen that was present in the atmosphere of the anaerobic chamber, and the bottles were recapped. CF and DCM were added as water saturated solutions (∼67 and 235 mM, respectively). After adding CF and/or DCM, the serum bottles were mixed on a shaker table (100 rpm for 1 h) to allow the CF and/or DCM to equilibrate between the headspace and liquid, sampled for analysis of the volatile compounds, and returned to the anaerobic chamber where they were incubated quiescently until the next sampling event. Incubation was at room temperature (∼22-24 ⁰C).

### Fate of CF using ^14^C-CF

The products from anaerobic degradation of CF were assessed using ^14^C-CF. Treatment #1 (Table 1) was performed in triplicate on three occasions (9 bottles total), with lactate as the electron donor. Degradation was allowed to continue until CF and DCM were below detection. For two of the sets of triplicates (6 bottles), the ability of the culture to consume unlabeled CF was confirmed prior to adding the ^14^C-CF. Treatment #2 was prepared identically but incubation was stopped when at least one third of the maximum possible stoichiometric level of DCM remained. Treatment #3 was the same as #1 except that BES (55 mM) was added to inhibit methanogenesis (23); two sets of triplicates were evaluated (6 bottles total). Treatment #4 served as a control to assess abiotic loss of ^14^C-CF with only medium A present.

CF-saturated water (0.25 mL/bottle) and a stock solution of sodium lactate (24 μL/bottle of a stock solution containing 650 g/L of 60% sodium lactate syrup) were added. The ratio of electron equivalents of lactate added to equivalents needed to reduce CF to DCM was approximately 20. Next, ^14^C-CF was added using an aqueous stock solution (see below). Headspace and liquid samples were used to establish the initial amount of ^14^C-CF present. Thereafter, the bottles were incubated quiescently. Headspace samples were removed periodically to determine the total levels of CF, DCM, CM, and methane by gas chromatography (see below). For treatment #2, triplicate bottles were sacrificed to determine the distribution of ^14^C (see below) when DCM degradation was incomplete. Treatments #1 and #3 were incubated until CF and DCM were no longer detectable, at which point these and the medium controls (treatment #4) were sacrificed to determine the distribution of ^14^C.

The presence of acetate at the start of the incubation was a consequence of prior additions of lactate to the culture. Acetate is a product of lactate fermentation. The initial concentration of acetate was ∼27 mM. Prior evaluation of the KB-1^®^ culture (from which KB-1^®^ Plus was developed) indicates that acetate does not serve as an electron donor for reductive dechlorination (24).

### Evaluation of CF and DCM as sole substrates

Treatments #5-10 explored the potential for CF and DCM to serve as sole substates for the KB-1^®^ Plus culture; ^14^C-CF was not used. Treatment #5 received DCM as the source of carbon and energy. After an initial dose of 16.5 µmol DCM per bottle was consumed, a second dose of 165 µmol was added. Treatment #6 was identical to #5 except that BES was added, along with one dose of DCM (16.5 µmol was consumed). Methane and hydrogen were quantified along with DCM.

Treatment #7 received CF and DCM. After the initial amount of CF was consumed (1.68 µmol per bottle), increasing amounts were added (up to 98 µmol), while the amount of DCM added remained constant (∼15 µmol per bottle). Prior to setting up the bottles for treatments #5-7, DCM (165 µM) was added to one liter of the KB-1^®^ Plus culture to confirm the activity of the culture. As with treatment #1, 2, 3, 5, and 6, acetate was present at the start of the incubation as a product of prior addition of lactate as the electron donor.

Treatment #8 differed from #7 as follows: The source of inoculum was one of the serum bottles from treatment #7 (100 mL); the culture was washed to remove residual organic acids; the initial dose of CF was lower (4.0 µmol per bottle) and a subsequent CF dose added was higher (16 µmol per bottle); and three lower doses of DCM were added (0.33, 0.16, and 3.47 µmol per bottle on days 0, 14, and 56, respectively). Washing was accomplished by centrifugation (1880-*g*, 30 min), decanting, and resuspending in 145 mL of medium B. Acetate was below detection in the resuspended culture. After washing, the color of resazurin in the medium turned slightly pink, indicative of a redox level above -110 mV. To restore a lower redox level, 8.8 µmol of Na_2_S was added (100 µL of a stock solution of 88 mM Na_2_S·9H_2_O). The washed culture was dispensed to triplicate 70 mL serum bottles, each with 44 mL of culture.

Two controls accompanied treatment #8. Treatment #9 consisted of the washed and resuspended KB-1^®^ Plus culture, with no CF or DCM added; 3.0 mL was dispensed to triplicate 12 mL serum bottles. Treatment #10 consisted of only medium B in 70 mL serum bottles, with similar initial doses of CF and DCM. At the end of the incubation period, aliquots from treatments #8-10 were analyzed for organic acids.

The pH in treatments #7 and 8 was monitored periodically; when the pH fell below 6.5 in treatment #7, sodium bicarbonate (100 µL of a stock solution of 10 g/L sodium bicarbonate) was added (day 398) to restore circumneutral conditions. The pH remained circumneutral in treatment #8.

### Analytical methods

^14^C-CF was purified as previously described for ^14^C-labeled trichloroethene (25). Briefly, the ^14^C-CF stock solution (100 μL) was injected onto a gas chromatograph and passed through the same packed column used to analyze the volatile organic compounds (see below). The outlet of the column was connected to stainless steel tubing that exited the GC oven and terminated with a needle, which was inserted into the microcosm headspace during the interval when CF eluted (∼6.5-7.0 min). N_2_ was used as the carrier gas (28.5 mL/min at 150 °C). Preliminary experiments with water controls confirmed that the only volatile compound present following this procedure was CF.

The total initial amount of ^14^C present in the serum bottles was quantified by counting samples of the headspace (0.5 mL) and liquid (1 mL) in 15 mL of liquid scintillation cocktail. The procedure to determine the distribution of ^14^C-labeled materials remaining in a microcosm at the time it was sacrificed is described by Darlington et al. (26). Volatile ^14^C-labeled biodegradation products (primarily DCM) were analyzed with a GC-combustion technique. A 0.5-mL headspace sample from the microcosms was injected onto the GC column and the separated compounds were routed to a catalytic combustion tube (containing CuO) at 800°C, where compounds were oxidized to CO_2_. Each fraction was then trapped in 3 mL of 0.5 M NaOH, which was then mixed with 15 mL of LSC. The efficiency of the combustion technique (i.e., Σ(^14^C in the fractions) divided by the ^14^C in a headspace sample) averaged 101±20%.

To measure ^14^CO_2_ and ^14^C-labeled nonvolatile products, liquid samples from the serum bottles were acidified and then sparged with N_2_, which was passed through an alkaline solution. The compounds remaining in the acidified sample were classified as nonstrippable residue (NSR); the activity in the alkaline solution was presumed to be ^14^CO_2_ (26). Formation of ^14^CO_2_ was confirmed by a precipitation test using barium hydroxide (26). The distribution of ^14^C includes an “unaccounted for” category, calculated as the total ^14^C remaining in a bottle minus the identified products (^14^C-CF, ^14^C-DCM, ^14^CO_2_ and ^14^C-NSR), and losses. Losses were calculated as the ^14^C added at time zero minus the total ^14^C remaining at the end of the incubation period.

The percentage of ^14^C-NSR in bottles for treatment #3 was sufficiently high (∼5% of the total ^14^C added) to warrant further characterization by fractionation on the same high performance liquid chromatograph (HPLC) used to quantify organic acids (see below). A filtered (0.20 µm) aliquot of NSR (100 µL) was injected onto the HPLC and fractions were captured in LSC as they eluted off the column, in intervals of 1.70 min (except for the intervals corresponding to formate and acetate, which were narrowed to 1.05 and 1.15 min, respectively). This was repeated with two more injections per bottle, so that the total volume of eluant captured per LSC vial was 1.9-3.1 mL. The ^14^C activity in each fraction was then counted (see below).

A PerkinElmer Tri-Carb^®^ 2910 TR liquid scintillation counter was used to quantify ^14^C activity. Corrections for counting efficiency were made according to a quench curve (sample spectral Quench Parameter-Ethernal (i.e., SQP(E)) versus efficiency) after incubating samples overnight in the dark.

Volatile organics (CT, CF, DCM, CM, and methane) were monitored by gas chromatographic analysis of headspace samples (0.5 mL), as previously described (27). Hydrogen was monitored in headspace samples using a gas chromatograph with a thermal conductivity detector (28). Results are reported in µmol per bottle. Based on the ratio of liquid to headspace and Henry’s Law constants, the conversion factors to mg per liter in the aqueous phase are 1.105 for CF, 0.08102 for DCM, and 0.4248 for CM. For hydrogen, the conversion factor from µmol per bottle to µM in the aqueous phase is 0.3062.

Organic acids (lactate, acetate, formate, citrate, and propionate) were measured in filtered (0.2 µm) samples (100 μL) by HPLC using a 3000 Ultimate Dionex HPLC system and an Aminex^®^ HPX-87H ion exclusion column (300-mm × 7.8-mm; BioRad). Eluent (5 mM H_2_SO_4_) was pumped (0.6 mL/min) through the column into a UV/Vis detector set at 210 nm. The minimum detection level was approximately 25 μM.

## Data Availability

Results that are not included in the manuscript may be found in the Supplemental Material.

## Supplemental Material

**Figures S1-S3:** replicates for Figure 1a;

**Figures S4-S6:** VOC results for treatment #2;

**Figures S7-S9:** replicates for Figure 1b;

**Figure S10:** VOC results for treatment #4;

**Figure S11:** replicates for Figure 3a;

**Figure S12:** replicates for Figure 3b;

**Figure S13:** replicates for Figure 4a;

**Figure S14:** replicates for Figure 4b; and

**Figure S15:** VOC results for treatment #10.

